# Improving the annotation of the cattle genome by annotating transcription start sites in a diverse set of tissues and populations using CAGE sequencing

**DOI:** 10.1101/2023.02.27.530265

**Authors:** M. Salavati, R. Clark, D. Becker, C. Kühn, G. Plastow, S. Dupont, G.C.M. Moreira, C. Charlier, E.L. Clark, the BovReg consortium

## Abstract

Understanding the genomic control of tissue-specific gene expression and regulation can help to inform the application of genomic technologies in farm animal breeding programmes. The fine mapping of promoters (transcription start sites [TSS]) and enhancers (divergent amplifying segments of the genome local to TSS) in different populations of cattle across a wide diversity of tissues provides information to locate and understand the genomic drivers of breed- and tissue-specific phenotypes. To this aim we used Cap Analysis Gene Expression (CAGE) sequencing to define TSS and their co-expressed short-range enhancers (<1kb) in the ARS-UCD1.2_Btau5.0.1Y reference genome (1000bulls run9) and analysed tissue- and population specificity of expressed promoters. We identified 51,295 TSS and 2,328 TSS-Enhancer regions shared across the three populations (Holstein, Charolais x Holstein and Kinsella beef composite [KC]). In addition, we performed a comparative analysis of our cattle dataset with available data for seven other species to identify TSS and TSS-Enhancers that are specific to cattle. The CAGE dataset will be combined with other transcriptomic information for the same tissues generated in the BovReg project to create a new high-resolution map of transcript diversity across tissues and populations in cattle. Here we provide the CAGE dataset and annotation tracks for TSS and TSS Enhancers in the cattle genome. This new annotation information will improve our understanding of the drivers of gene expression and regulation in cattle and help to inform the application of genomic technologies in breeding programmes.

## Introduction

The reference genome for domestic cattle, ARS-UCD1.2, now has a very high-quality annotation of both expressed and regulatory regions generated for the Hereford breed e.g. (Goszczynski et al, 2021; Halstead et al. 2020). There is, however, still very little available information about how the genome is expressed and regulated across different populations of domestic cattle. This lack of knowledge hinders efforts to define and predict the effects of genetic variants and link genotype to phenotype. To address this knowledge gap transcriptomic resources that include both multiple different tissue types and populations of cattle are required.

High resolution mapping of the actively transcribed regions of the genome can help to identify the drivers of gene expression, regulation and phenotypic variation (Tippens et al. 2018). Defining transcription start sites (TSS) within promoter regions provides information about how genes controlling traits of interest are expressed and regulated. Recently, the theory of multiple expression clusters within promoters has been used to annotate and fine map TSS within mammalian transcriptomes (Frith et al. 2008; Andersson et al. 2014). These putative core promoter and associated enhancer regions are defined using 5’ cap transcript sequencing e.g. via RAMPAGE (RNA Annotation and Mapping of Promoters for the Analysis of Gene Expression) (Batut and Gingeras 2013; Goszczynski et al. 2021) and CAGE (Cap Analysis Gene Expression) (Forrest et al. 2014; Robert et al. 2015; Deviatiiarov et al. 2017; Noguchi et al. 2017; Salavati et al. 2020; Ross et al. 2022). The fine mapping of promoters and enhancers, in this way, in different populations of farmed animals, across a wide diversity of tissues, provides information to link genotype to phenotype, by locating and understand the genomic drivers of breed- and tissue-specific phenotypes.

To improve TSS and enhancer annotation of the current reference genome for cattle (ARS-UCD1.2), we employed a diverse sampling approach to include transcriptomes from both sexes and multiple age categories (calves < 4 weeks, peri-puberty juveniles ∼ 7-8 months and adults > 1.5 years) from 3 divergent populations of cattle: Dairy (Belgian Holstein Friesian), Beef-dairy cross (German Charolais X Holstein F2) and Canadian Kinsella cattle (beef composite). Capture of gene promoters from different cattle breeds is important in identifying functional genomic features impacting selected or adapted traits in both dairy and beef populations (Halstead et al. 2020; Alexandre et al. 2021). Defining robust genomic annotations has proven to be useful in the sustained genetic improvement of farmed animals (Georges et al. 2018).

To this aim we used CAGE sequencing to define TSS and their co-expressed short-range enhancers (<1kb) (TSS-Enhancers) in the ARS-UCD1.2_Btau5.0.1Y reference genome (1000bulls run9) (Hayes and Daetwyler 2019) and analysed tissue- and population specificity of expressed promoters. We also utilised publicly available CAGE datasets (Forrest et al. 2014) for human, chicken, mouse, rat, macaque monkey and dog from the FANTOM5 project, and for sheep (Salavati et al. 2020), to provide a cross-species comparative analysis of TSS and TSS-Enhancers. Using comparative analysis this study provides a cattle-specific set of TSS and TSS-Enhancers in multiple tissues from dairy (Belgian Holstein), beef-dairy cross (Charolais x Holstein) and multi-breed composite beef (KC) cattle. Several transcriptomic datasets (RNA-Seq and small RNA-Seq) are being generated from the same set of tissues, as part of a wider effort in the BovReg project, to generate a high resolution transcriptomic map to improve the annotation of the ARS-UCD1.2 reference assembly, by adding transcriptomic information for multiple tissue samples across the three different populations. Additional annotation information will improve our understanding of the drivers of gene expression and promoter diversity/plasticity in cattle and help to inform the application of genomic technologies in breeding programmes.

## Materials & Methods

### Animals

Samples from three diverse cattle populations were chosen for the purpose of this study: Dairy (Holstein Friesian), beef x dairy (Charolais x Holstein F2) and composite beef (Kinsella composite [KC; Angus, Hereford and Gelbvieh breeds account for approx. 65% of the breed composition of the samples with signals from 9 other cattle breeds including Brown Swiss, Limousin, Simmental, Holstein and Jersey]) lineages. Tissues were collected from two animals (1 male and 1 female per population = 6 animals in total) from each population. These 6 animals included three different age groups: Holstein Friesian calves from Belgium (neonatal: male calf 24 days and female calf 22 days), KC steer (bullock 217 days) and heifer (juvenile, 210 days) from Canada and Charolais x Holstein F2 cow and bull (adult: bull 18 months and cow 3 years, 7months and 13days) from Germany. Necropsy and tissue collections were performed under site-specific ethics approval by qualified research personnel at University of Alberta Canada (Animal Use Protocol #00002592), University of Liege, Belgium (*Commission d’Etique Animale; Dossier* #17-1948) and the Research Institute for Farm Animal Biology, Germany. In Germany, all experimental procedures were performed according to the German animal care guidelines and were approved and supervised by the relevant authorities of the State Mecklenburg-Vorpommern, Germany (State Office for Agriculture, Food Safety and Fishery; LALLF M-V/ TSD/7221.3–2.1-010/03).

### Sample collection

A total of 102 samples from 24 different tissues were collected from the 6 animals (3 populations, different ages and 2 sexes). Tissue representation for each population was as follows: dairy (Holstein, n=43 tissues), beef x dairy cross (Charolais x Holstein, n=31 tissues) and composite beef (KC, n=31 tissues). Details of the collected tissues are shown in Table 1. Tissue samples were snap frozen immediately upon collection, stored at -80°C for downstream RNA extraction and for the beef x dairy cross and composite beef samples shipped on dry ice to a central location (GIGA, University of Liège, Belgium) for RNA isolation.

**Table 1.** List of all the samples collected and sequenced by CAGE-Seq including 24 tissue types, from 6 animals (3 populations, 3 ages and 2 sexes). Belgian Holstein Friesian: HF, German Charolais x Holstein F2: Char x Hol, Canadian Kinsella composite: KC.

| Tissue | Male calf HF | Female calf HF | Bull Char x Hol F2 | Cow Char x Hol F2 | Steer/bullock KC | Heifer KC |
| --- | --- | --- | --- | --- | --- | --- |
| Adrenal Gland Cortex | X | X | X | X | X | - |
| Cerebellum | X | X | X | - | - | X |
| Cerebrum Cortex | X | X | X | X | X | - |
| Colon | X | X | X | X | X | X |
| Duodenum | X | X | X | X | X | X |
| Heart | X | X | X | - | X | X |
| Hypothalamus | X | X | - | - | - | - |
| Ileum | X | X | X | X | X | X |
| Jejunum | X | X | X | X | X | X |
| Kidney | X | - | X | X | X | X |
| Liver | X | X | X | X | X | X |
| Lung | X | - | X | X | X | X |
| Lymph Node | X | X | X | X | X | X |
| Mammary Gland | - | X | - | X | - | X |
| Ovary | - | X | - | - | - | X |
| Pancreas | X | X | - | - | - | - |
| Pituitary Gland | - | X | - | - | X | X |
| Rumen | X | X | X | X | X | X |
| Skeletal Muscle | X | X | - | - | - | - |
| Spleen | X | X | X | X | X | X |
| Subcutaneous Fat | X | X | - | - | - | - |
| Testis | X | - | X | - | - | - |
| Thyroid Gland | X | X | X | X | - | - |
| Uterus | - | X | - | X | - | X |

### RNA extraction and quality control

Total RNA was extracted using miRNeasy kit (QIAGEN) from the snap-frozen tissues samples, following the protocol provided by the manufacturer for the purification of Total RNA from Animal Tissues. The RNA integrity (RIN) was detected by the Agilent Bioanalyzer system (Agilent Technologies, Santa Clara, CA, USA). Aliquots containing 5μg of total RNA (RIN >7) were then stored at -80° C before shipping to Edinburgh Clinical Research Facility, Edinburgh, UK.

### CAGE-Seq library preparation and sequencing

CAGE libraries were prepared from 5μg of total RNA (post DNase treatment) according to (Takahashi et al. 2012). A modification of the original barcodes from the Takahashi et al. (2012) protocol (3nt length) was required in order to perform sequencing on the Illumina NextSeq 550. This modification introduced 6nt length barcodes for multiplexing of the libraries. The original barcodes: ACG, GAT, CTT, ATG, GTA, GCC, TAG, and TGG were extended to a set of 21 unique 6nt barcodes. Overall 13 library pools were produced and sequenced on an Illumina NextSeq 550 (50nt single end as previously described in (Salavati et al. 2020) in 7 different runs. The details of the barcode assignments to each sample and the pool ids are described in Supplementary_file_1.xlsx.

### CAGE-Seq data analysis

The analysis pipeline was developed using NextFlow workflow scripting (di Tommaso et al. 2017). The pipeline was built using the previously described steps in https://bitbucket.org/msalavat/cagewrap_public/src/master/. After demultiplexing, trimming and quality control, the reads were mapped against the ARS-UCD1.2_Btau5.0.1Y assembly run 9 (Hayes and Daetwyler 2019) using the nf-cage pipeline (Salavati and Espinosa-Carrasco 2022 Jul 18). The base-pair resolution output bigWig files (2 files per sample +ve and -ve strand; n=204 for 102 samples) were loaded in RStudio (RStudio Team 2015) (R > v4.0.0) for downstream analysis using the CAGEfightR v1.16.0 package (Thodberg et al. 2019).

### Transcription start site and enhancer prediction analysis

The putative transcription start sites (TSS) and TSS-Enhancer regions were identified using the uni- and bi-directional clustering algorithms in CAGEfightR v1.16.0 as described in (Thodberg et al. 2019). Clustering overlapping same-strand CAGE tags mapped to either strands of the DNA was considered uni-directional, compared to clustering of non-overlapping tags mapped within 400-1000bp of each other to opposing strands (e.g. gene +ve with a nearby eRNA -ve or *vice versa*) using a bi-directional clustering approach. CAGE tag TSS clusters (CTSS) and their normalised expression profile (CTPM; CAGE tags-per-million mapped) were produced using quickTSS and quickEnhancers functions of the CAGEfightR package v1.16.0. For both TSS and TSS-Enhancer regions a minimum 10 reads per CTSS and 2/3^rd^ sample support (i.e. if the CTSS was present in a minimum of 66/102 tissues) were imposed as filtration criteria, as previously described in (Salavati et al. 2020). The putative regions were annotated using the assignTxID, assignTxType, assignGeneID and assignMissingID functions of the CAGEfightR v1.16.0. The Txdb object used for annotating the CAGE-Seq dataset was built using the *<u>Bos_taurus</u>*.*<u>ARS-UCD1</u>*.*<u>2</u>*.*<u>106</u>*.*<u>gff3</u>*.*<u>gz</u>* file from Ensembl v106.

### Mapping significant TSS and TSS-Enhancer co-expression links

Co-expression of the predicted TSS and TSS-Enhancer regions was tested using a Kendall correlation test (*p*<0.05 sig. followed by Benjamini-Hochburg adjustment; FDR < 0.01). The co-expressed pairs were identified using the findLinks function of the CAGEfightR v1.16.0 as previously described (Thodberg et al. 2019; Thodberg and Sandelin 2019) and annotated using the Bos_taurus.ARS-UCD1.2 Ensembl v106 gene models. Using the gap (in bp) between the TSS (query) and Enhancer (subject) and the assigned gene symbol to either region, 3 groups of links were created: *cis* [same gene] where TSS and Enhancer regions had a gap less than 1kb, *trans* [nearby gene] where the gap was larger than 1kb and *novel* (cis or trans) where there was no gene annotation available for either of the linked pair. The gap size (in bp) and the Kendal correlation coefficient (range = [-1,1]) of this co-expression analysis was then used for further investigation of these links. A two dimensional kernel density estimate was calculated for the gap between linked TSS and Enhancers versus the link’s correlation coefficient. This analysis was performed using the MASS package v7.3-58.1 (Venables and Ripley 2002) (MASS::kde2d) and visualised using ggplot2 v 3.3.6 (Wickham 2009) (ggplot2::geom_density2d_filled) in R.

### Identification of long range enhancer stretches present in the cattle genome

A hierarchical clustering of the TSS-Enhancer regions (obtained using the bi-directional analysis method in the CAGEfightR package) was performed to identify any super-enhancers. A 10kb window scan was performed to locate stretches of the genome containing at least 3 Enhancers within a window. This analysis was performed using the findStretches function of the CAGEfightR v1.16.0 followed by a Kendal correlation test of the expression matrix (CTPM values as input).

Three genomic regions harbouring copy number variants associated with milk traits (CNV6 [chr13:70,496,054-70,623,303], CNV28 [chr7:42,700,425-42,788,788], and CNV33 [chr17:73,055,503-75,058,715]) within the cattle genome (UMD3.1), previously reported by Xu et al. (Xu et al. 2014), were lifted over to the ARS-UCD1.2 coordinates using the UCSC liftover tool (Hinrichs et al. 2006). The super-enhancer stretches identified in the cattle CAGE dataset were overlaid with the lifted over CNV regions using IGVtools (Robinson et al. 2011; Thorvaldsdóttir et al. 2013)

### Characterising tissue-specific TSS and TSS-Enhancers

Tissue specific sets of TSS and TSS-Enhancers were produced in 24 separate runs of the 2 clustering algorithms (quickTSS and quickEnhancers). All samples of the same tissue type were used to create tissue specific outputs (min 10 reads/CTSS and support 2≤n≤6). The tissue (Raivo Kolde) specific TSS and TSS-Enhancer regions were also annotated using the Ensembl v106 gene models as described in the TSS and Enhancer prediction analysis section. The expression matrix (CTPM) of all identified TSS across all tissue types was used to produce a heat map based on tissue specificity indexes (TSI ranging from 0 = no expression in a particular tissue to 1 = only expressed in a particular tissue). The TSI indexes for each TSS were produced using tspex v0.6.1 (Camargo et al. 2020 Aug 4) and visualised using pheatmap v1.0.12 (Julien et al. 2012) in R.

### Characterising population specific TSS and TSS-Enhancers in the cattle dataset

Population specific sets of TSS and TSS-Enhancers were analysed by applying the uni- and bi-directional clustering algorithms three times to all tissue samples from each population of cattle: 2 Holsteins (41 samples), 2 Charolais x Holstein F2s (31 samples) and 2 KC composite (30 samples). In each run only TSS and TSS-Enhancers present in all tissue types (100% support) were kept for further analysis i.e. to define a TSS or TSS-Enhancer as Holstein specific it had to be present in all Holstein derived samples. A Holstein signature of TSS and TSS-Enhancers (based on start-end coordinates) was established as follows: Firstly a set of TSS and TSS-Enhancer regions present in all 3 population sets (CHAR:KC:HOL_signature) was created, then a set shared only between Holstein Friesian and Charolais x Holstein F2 sets (CHAR:HOL_signature) was created and finally a set shared only between Holstein Friesian and Kinsella composite sets (KC:HOL_signature) was created. An intersection analysis was then performed using UpSetR v1.4.0 (Lex et al. 2014).

### Comparative analysis using the Fantom5 and sheep CAGE datasets

Mapped CAGE datasets from human (hg19, n = 152), rat (rn6, n=13), mouse (mm9. n=17), chicken (galGal5, n=32), dog (canFam3, n= 13) and Macaque monkey (rheMac8, n=15) were obtained from (Bertin et al. 2017). The CAGE dataset for sheep (PRJEB34864) (Salavati et al. 2020) was re-analysed by mapping against the ARS-UI_Ramb_v2.0 (GCF_016772045.1) reference genome from NCBI v106. After re-mapping of these 56 ovine tissue samples, the TSS regions were annotated using the CAGEfightR v1.16.0 and GCF_016772045.1_ARS-UI_Ramb_v2.0_genomic.gff.gz gene models. The identified TSS regions and their annotated gene symbols (i.e. Ensembl attribute GENE NAME and NCBI RefSeq GENE SYMBOL) were extracted from each of the datasets for comparative analysis. TSS regions were annotated by gene symbols in all 8 datasets (in sheep and cattle using CAGEfightR assignGeneID plugin). Then merged based on sharing the same gene symbol (i.e. homologues) or not to form 5 groups: Avian/Mammalian homologues for TSSs present in all 8 species datasets, Mammalian specific TSS found in all 7 mammalian species, Human specific for TSS present only in human and species specific for all other uniquely assigned TSS. This analysis reduced the number of TSS in each dataset to only those with a gene symbol annotation nearby. The majority of species specific TSS for each dataset had either a unique gene symbol or were novel genes followed by unannotated TSS regions.

### Statistical analysis and data visualisation

All statistical analysis and data visualisations were carried out in R > v4.0.0 using RStudio (RStudio Team 2015) and tidyverse suite v1.3.2 (Wickham et al. 2019). The nf-cage pipeline was run on the high performance computing cluster of the University of Edinburgh (Eddie) (Edinburgh 2020).

## Results

### CAGE-Seq library size and mapping metrics

An average (± SE) of 15.5±0.53 million reads per CAGE sample were generated. After mapping to the ARS-UCD1.2_Btau5.0.1Y (Hayes and Daetwyler 2019) reference genome a 94% average mapping rate was achieved for all of the tissues (24 types) within the dataset (n=102).

### CAGE-Seq initial clustering and quality control

After Initial CAGE tag clustering (CTSS) more than 4.3 million putative TSS (uni-directional) and 57,078 TSS-Enhancer (bi-directional) regions were identified in total. A minimum of 10 reads per region was the only filtering criteria set at this stage of the analysis, with the 2/3rds rule being applied later. The tissue grouping of the TSS and TSS-Enhancer regions is shown in Figure 1.

**Figure 1.**
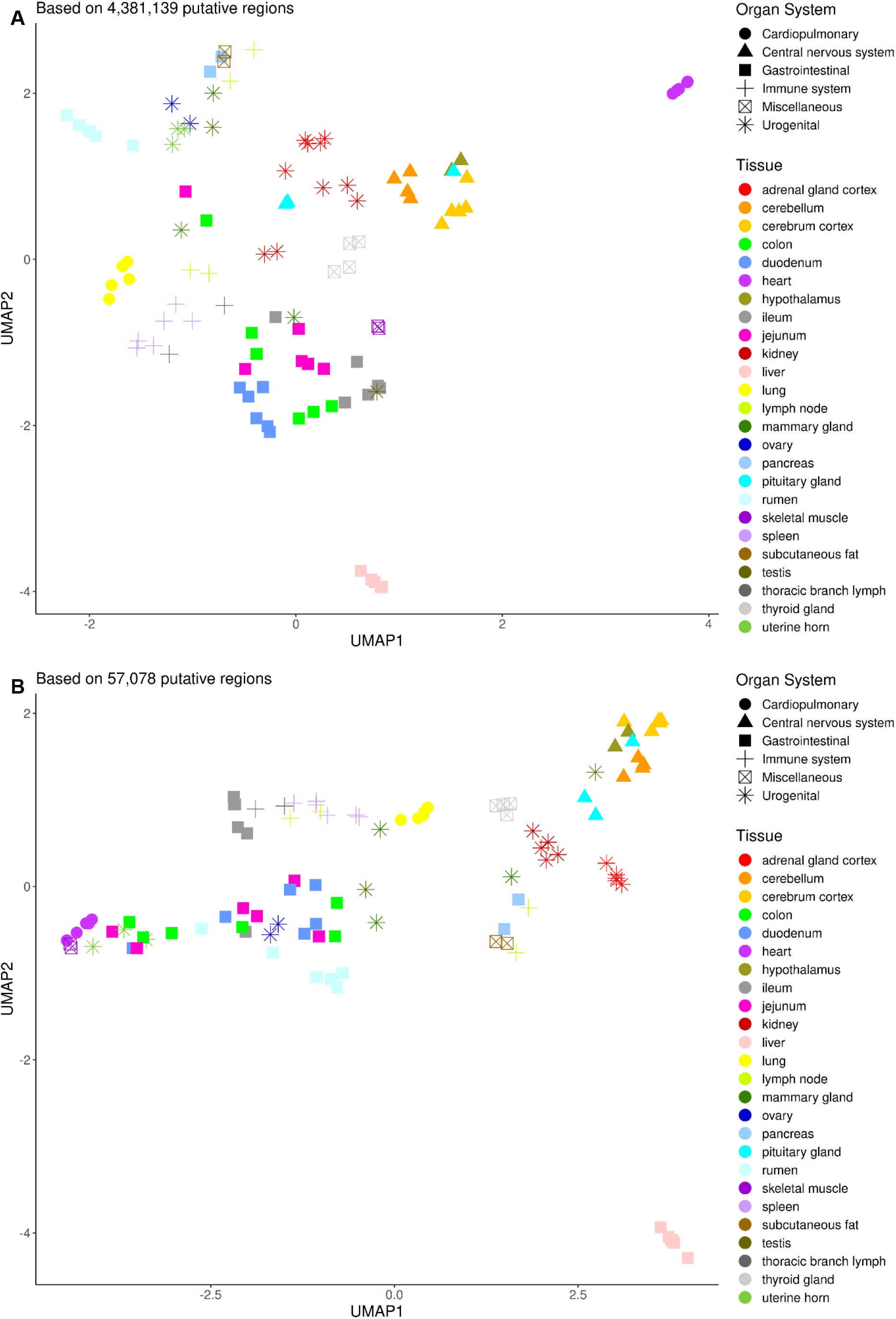
Dimension reduction of the cattle CAGE-seq dataset using uniform manifold approximation and projection (UMAP). A) The putative TSSs (4,381,139 regions of the cattle genome) and their expression values (CTPM) for all the 102 tissue samples were used as the input matrix for UMAP. The first 2 components are visualised with tissue name (colour) and organ systems (shapes) as labels. B) The putative TSS-Enhancers (57,078 regions of the cattle genome) and the respective CTPM values were used as the input matrix for UMAP. The first 2 components are visualised with tissue name (colour) and organ systems (shapes) as labels.

The gastrointestinal (GI) tract tissues (shown as squares in Figure 1A) and immune system tissues (lymph nodes and spleen indicated by a + sign in Figure 1A) formed relatively distinct clusters as expected. Although this grouping was less pronounced in the TSS-Enhancer profiles for the immune system tissues, the GI tissues kept the original grouping structure, as shown in Figure 1B. Specific tissues e.g. rumen, liver and heart were clustered very distinctly and consistently across TSS and TSS-Enhancers profiles.

### Identifying pervasive TSS and TSS-Enhancers across tissues

We considered a putative TSS or TSS-Enhancer region, real/reproducible only when it was present across at least 2/3^rds^ of the tissues. After filtering using the 2/3^rds^ rule 51,295 TSS and 2,328 TSS-Enhancers were detected for cattle with a mean of 91 ± 0.04 (median 94) samples supporting each putative region. This sample support translated into mean 23.7 ± 0.002 (median 24) tissue-type support for each region. Similar metrics for the sheep dataset (PRJEB34864) were captured after remapping and applying the same tissue representation criteria (Table 2). Overall 15,364 genes and 27,588 corresponding transcripts were annotated using the CAGE dataset we generated for cattle. We identified 51,295 TSS regions of which 16,957 (33%) were novel and 34,338 overlapped current gene models (Ensembl v106). From the novel putative TSS regions more than 2/3^rds^ (67%) resided within intergenic coordinates from the ARS-UCD1.2 gene build models (Ensembl v106) and 5,592 mapped to antisense features.

**Table 2.** Mapped and annotated CAGE-Seq uni-directional clusters (TSS regions) in sheep mapped to (ARS-UI_Ramb_v2.0) and cattle mapped to (ARS-UCD1.2_Btau5.0.1Y) using reference assembly gene models (using the min 2/3^rd^ tissue representation threshold).

| Genomic Region | Sheep - ARS-UI_Ramb_v2.0 |  |  |  | Cattle - ARS-UCD1.2_Btau5.0.1Y |  |  |  |
| --- | --- | --- | --- | --- | --- | --- | --- | --- |
| | Novel | Anno<sup>*</sup> | Total | %<sup>\$</sup> | Novel | Anno<sup>*</sup> | Total | %<sup>\$</sup> |
| Promoter | 0 | 13,372 | 13,372 | 39.4 | 0 | 9,763 | 9,763 | 19 |
| proximal | 0 | 944 | 944 | 6 | 0 | 3,296 | 3,296 | 6.4 |
| fiveUTR | 0 | 873 | 873 | 6.7 | 0 | 2,975 | 2,975 | 5.8 |
| threeUTR | 0 | 2,197 | 2,197 | 7.9 | 0 | 2,118 | 2,118 | 4.1 |
| CDS | 0 | 4,513 | 4,513 | 16.1 | 0 | 8,355 | 8,355 | 16.3 |
| exon | 0 | 295 | 295 | 1.9 | 0 | 238 | 238 | 0.5 |
| intron | 0 | 2,386 | 2,386 | 10 | 0 | 7,593 | 7,593 | 14.8 |
| antisense | 1,034 | 0 | 1,034 | 4.2 | 5,592 | 0 | 5,592 | 10.9 |
| intergenic | 1,397 | 0 | 1,397 | 7.9 | 11,365 | 0 | 11,365 | 22.2 |
| Total TSS | 2,431 | 24,580 | **27,011** | 100 | 16,957 | 34,338 | **51,295** | 100 |
| Total TSS-En<sup>£</sup> | 34 | 1,459 | **1,493** |  | 373 | 1,955 | **2,328** |  |
| Annotated genes/transcripts |  |  | 13,771 / 45,298 |  |  |  | 15,364 / 27,588 |  |

\* Annotated using the reference assembly gff3 track
\$ Percentage calculated based on total per genomic region category / total TSS clusters
£ TSS-Enhancers (Bi-directional clustering)

The median number of putative TSS regions per gene and transcript model were 1 and 2 respectively (mean 1.6 TSS/gene and 3.1 TSS/transcript). All the identified TSS and TSS-regions were annotated using the current Ensembl v106 gene builds. The majority of the annotated regions resided within the promoter and/or 1kb proximal of the first exon. A larger portion of the TSS regions (22.2%) in the cattle dataset fell within intergenic (no gene annotation in ARS-UCD1.2 Ensembl gff3) coordinates compared to the sheep dataset (ARS-UI_Ramb_v2.0 NCBI gff3). The breakdown of the cattle CAGE dataset annotation based on genomic feature category is shown in Figure 2.

**Figure 2.**
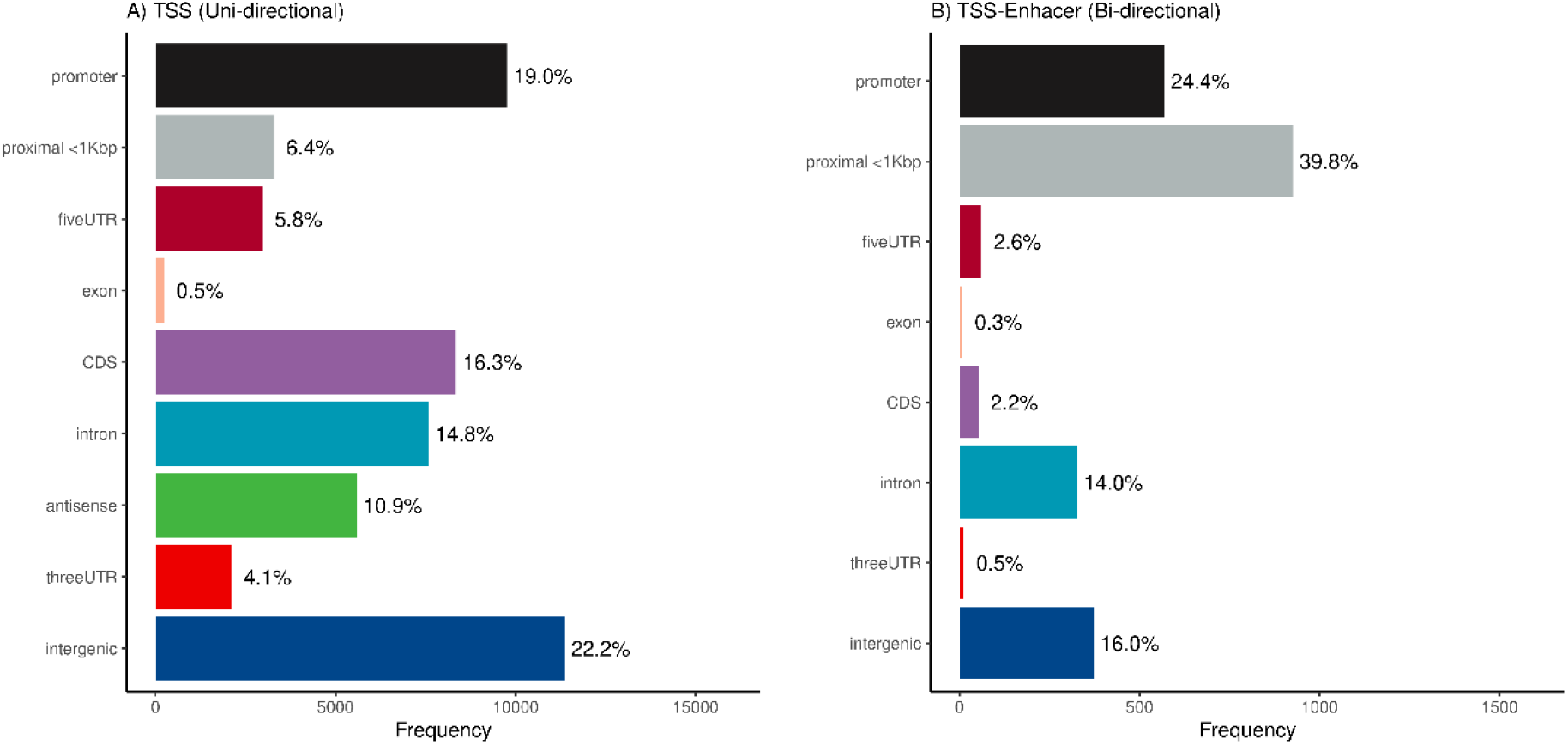
Genomic feature annotation of the cattle CAGE dataset based on the Ensembl v106 annotation. A) Frequency distribution of the putative TSS regions identified in at least 2/3^rd^ of the sampled tissues. B) Frequency distribution of the putative TSS-Enhancer regions identified in more than 2/3^rd^ of the sampled tissues.

### Identifying co-expressed TSS and Enhancers regions

We identified significant (Kendal correlation adjusted *p* < 0.01) co-expression between bi-directional clusters (TSS-Enhancer region) and multiple uni-directional clusters (TSS) in both the cattle and sheep CAGE datasets. After applying the 2/3rds of tissues representation threshold, an average 3.73±0.05 (median 3) TSS in sheep and 6.62±0.12 (median 5) TSS in cattle showed significant co-expression with a neighbouring Enhancer region. We identified 3,641 co-expression links in sheep and 15,600 in the cattle dataset. The average Kendall estimates of these significantly co-expressed links were 0.46 ± 0.003 and 0.34 ± 0.001 for the sheep and cattle tissues respectively. The expression patterns and correlation estimates for the cattle dataset are shown in Figure 3.

**Figure 3.**
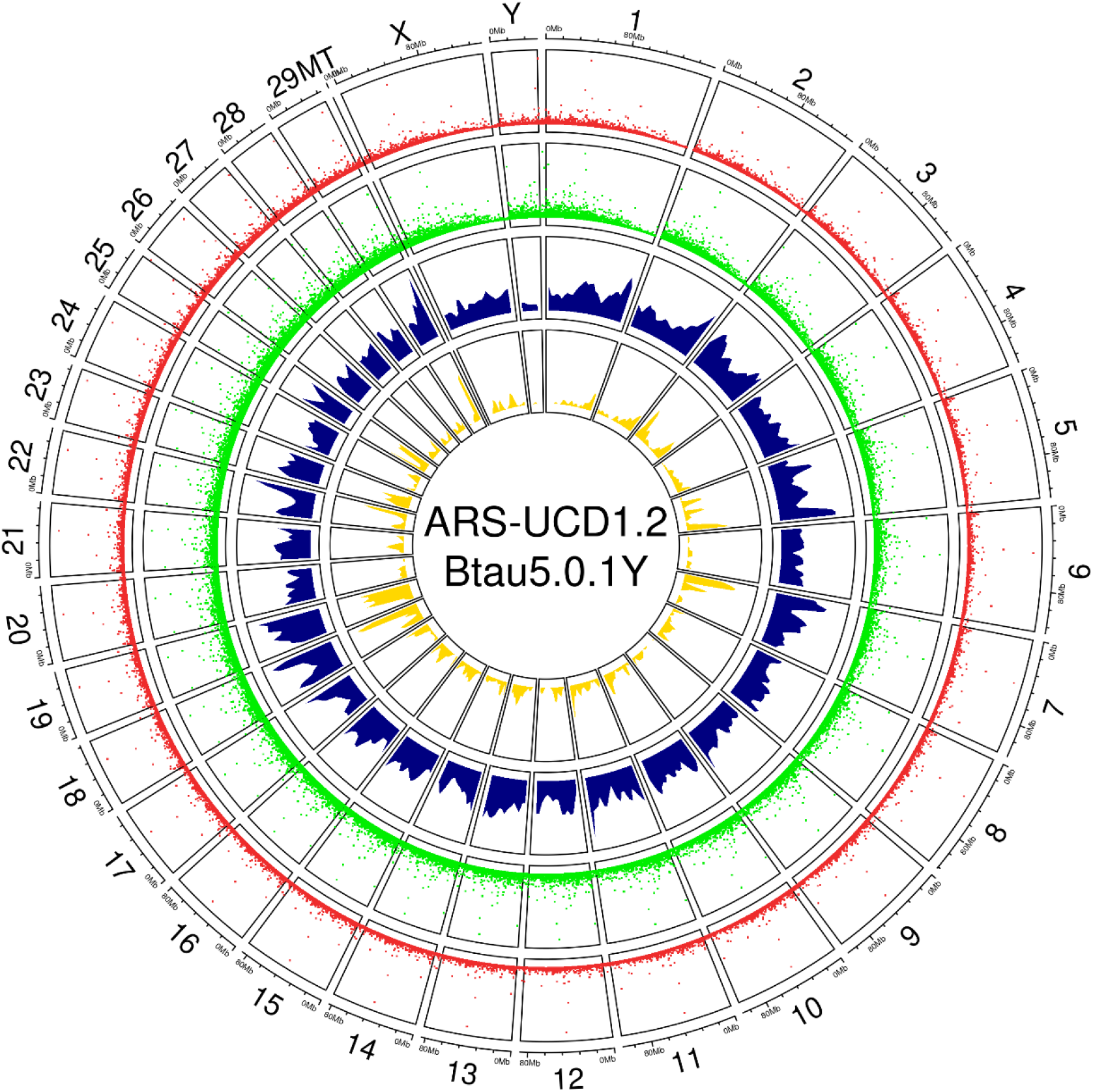
Distribution of uni-directional (TSS) and bi-directional (TSS-Enhancer) CAGE clusters within the cattle genome (ARS-UCD1.2_Btau5.0.1Y). The TSS clusters (red), TSS-Enhancer (green), significant positive (blue) and negative (yellow) correlation between co-expressed Enhancer and TSS(s) are shown in genomic tracks. The height of the tracks shows scaled expression or correlation coefficients (0-1).

We further analysed the co-expression of TSS and Enhancer regions using a 2 dimensional density map. The Kernel Density Estimate (KDE) was used to identify co-expression signals based on correlation estimates vs relative distance from TSS. These signals in both annotated and unannotated genomic coordinates of the cattle dataset have been visualised in Figure 4.

**Figure 4.**
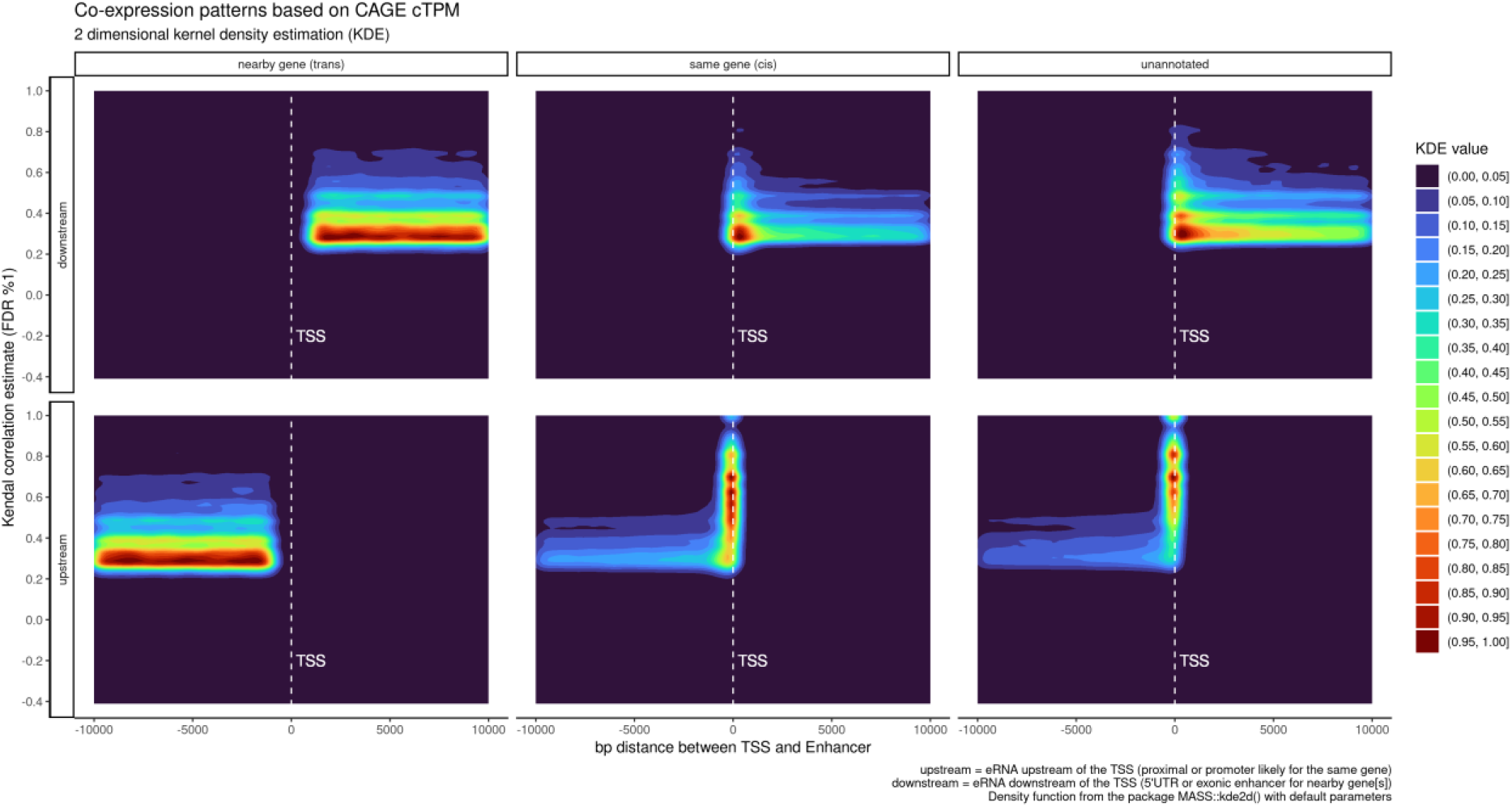
Kernel density estimates of correlation coefficient (0-1) and distance to TSS (bp) of all significant co-expression profiles within the cattle CAGE dataset. The Kendal correlation estimates and the distance between the Enhancer region and associated TSS were used in the KDE analysis. Enhancer activity within 1kb vicinity of the TSS was considered as the “same gene”, between 1kb-10kb “near by gene” while all unannotated putative TSS (termed ‘Novel’) were linked with annotated Enhancer regions marked as “unannotated”.

The KDE analysis showed a stronger co-expression (average estimate of 0.44; Welch test *p* < 0.01) for short range (< 1kb to TSS) in both upstream and downstream enhancer RNA (eRNA) compared to long range (average estimate 0.38). The longer genomic distance between TSS and co-expressed Enhancers was expected to result in smaller correlation estimates. The average (up- and downstream) co-expression correlation estimate of 0.38 was with nearby genes (1kb-10kb windows) pointing to this decay of co-expression due to the distance. Unannotated TSS and Enhancer links showed the highest average correlation estimates (0.47 Welch test *p* < 0.01) compared to the other 2 categories. Further details of the comparison between groups can be found in Supplementary Figure 1.

### Identifying long stretches of Enhancer activity in the cattle genome

The analysis of the ‘super enhancers’ (stretches of bi-directional CAGE clusters) encompassing multiple enhancers within each stretch in the sheep dataset resulted in 2 super enhancer predictions. These stretches were formed of 6 TSS-Enhancer clusters with the longest stretch of 5,172bp [cluster of 3 enhancers]. Similar analysis of the cattle CAGE dataset from 3 populations resulted in 16 super enhancer stretches from 53 TSS-Enhancer clusters. The longest stretch was 25,679bp which contained 6 TSS-Enhancers. The number of discovered super enhancer regions overall was higher in the cattle dataset (3 populations) compared to the sheep (which came from a single individual). The detail of the enhancer stretches and their coordinates can be found in supplementary File 2.zip.

We also overlaid the enhancer stretches with previously reported copy number variant (CNV) regions of the cattle genome associated with milk production traits in Holsteins (Xu et al. 2014) Three milk trait associated CNVs (chr7, ch13 and chr17 of UMD3.1 lifted to ARS-UCD1.2) had large overlaps with TSS-Enhancers identified in the following genes: *PLCG1* (CNV at chr13: 13:69,794,566-69,921,810), *PPM1F* (CNV at chr17:71,988,770-71,998,055), *TOP3B* (CNV at chr17:71,964,684-71,967,648) and *TANGO2* (CNV at chr17:72,965,809-72,970,736).

### Identifying tissue specific TSS and TSS-Enhancer regions

Tissue-specific analysis captured, on average 253,852 ± 24,713 (± SE) TSS clusters per tissue, 41.6% of which were novel. On average 12,138 ± 889 TSS-Enhancer clusters per tissue were captured (27.6% novel). Including multiple biological replicates per tissue type resulted in a higher number of genes being annotated by the cattle CAGE dataset compared to the Ensembl v106 reference annotation. We captured significantly (adjusted *p* < 0.05 Tucky HSD post ANOVA) less genes and transcripts annotated by CAGE tags in tissue types with 2-3 replicates compared to higher (n>4) biological replicates (Figure 5) Clustering of the tissues based on the tissue specificity index (TSI) (row wise transformed CTPM) (Figure 6) showed tissue-specific promoter activity present in testis, central nervous system tissues, gastrointestinal tract and tissues with a higher epithelial density of immune cells e.g. ileum, mammary gland, lungs, spleen and lymph nodes.

**Figure 5.**
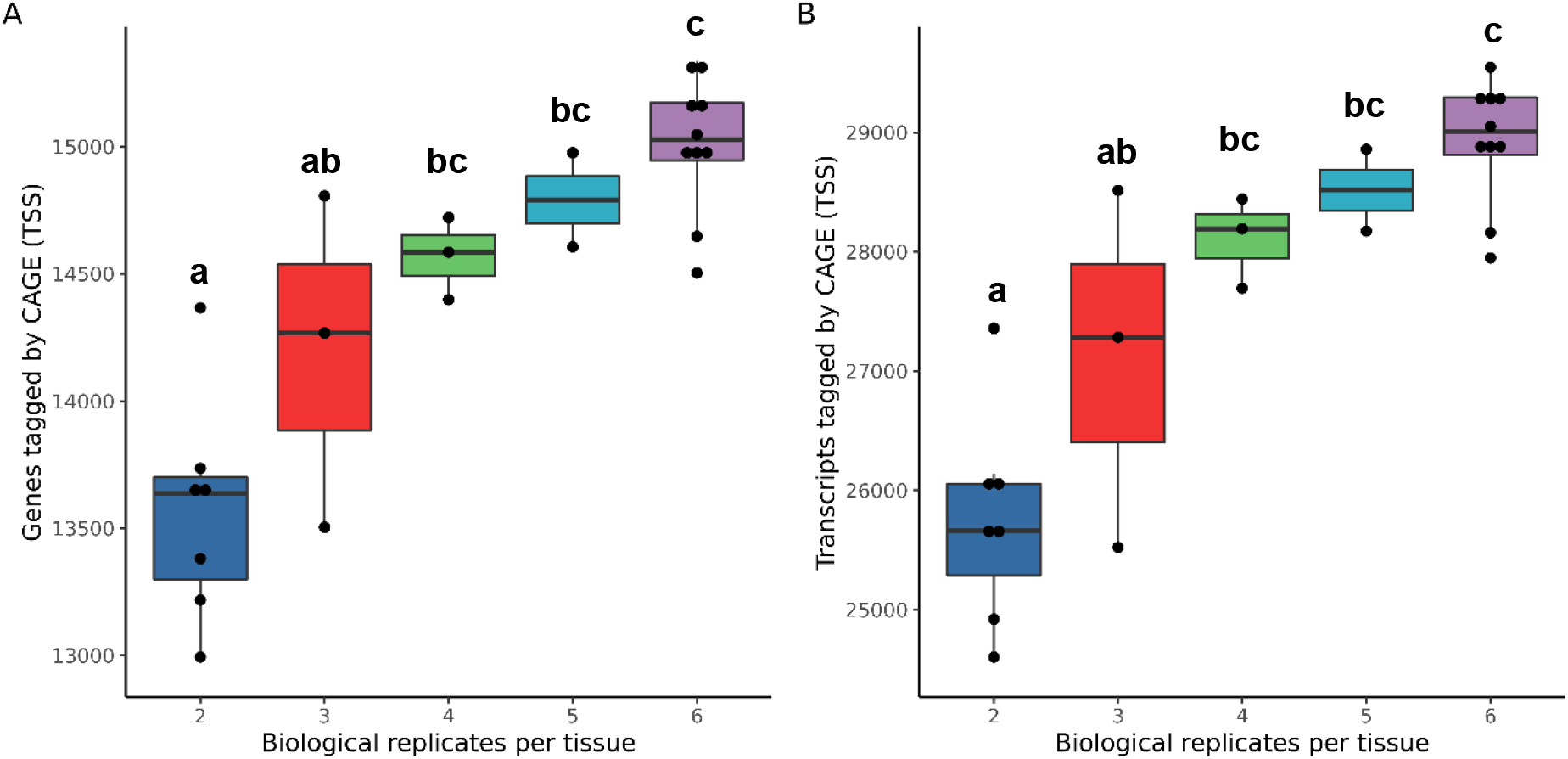
Number of replicates per tissue type and its effect on genes (A) and transcripts (B) annotated by the cattle CAGE dataset. All 24 tissue types were grouped by the number of biological replicates/samples previously described in Table 1. The significant difference between 5 groups was tested using ANOVA followed by stats::TukeyHSD in R. The significant adjusted p values are marked by letters “a”, “b” and “c”.

**Figure 6.**
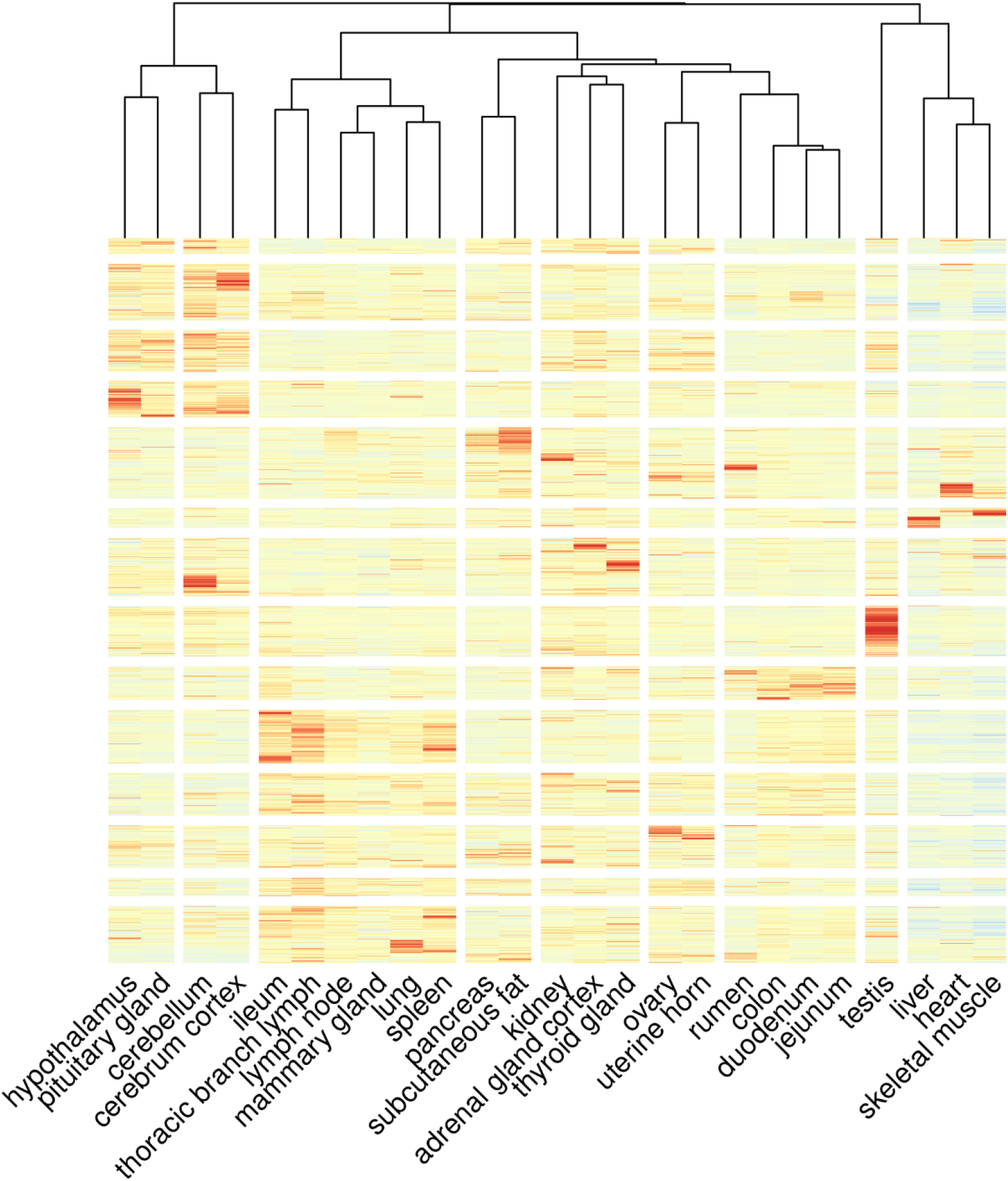
Tissue specificity indices (TSI) of the all the putative TSS regions (rows) based on CTPM and tissue type (columns). The heatmap was built using a row and column wise clustering algorithm (hclust ∼ Manhattan distances) and the averaged TSI of the TSS across tissue replicates.

### Population specific TSS and TSS-Enhancer regions

Population-specific analysis showed differences in TSS coordinates and expression levels between the 3 populations of cattle (Holstein Friesian [HOL], Charolais x Holstein and Kinsella composite [KC] beef cattle). The highest number of population-specific TSS were found in the KC-composite (3,120) followed by 1,140 in Holstein and 1,106 in Charolais x Holstein. The same pattern was observed in the TSS-Enhancer regions (414 in KC-composite, 281 in Charolais x Holstein and 202 in Holstein). The detailed population-specific sets of TSS and TSS-Enhancer regions are shown in Figure 7.

**Figure 7.**
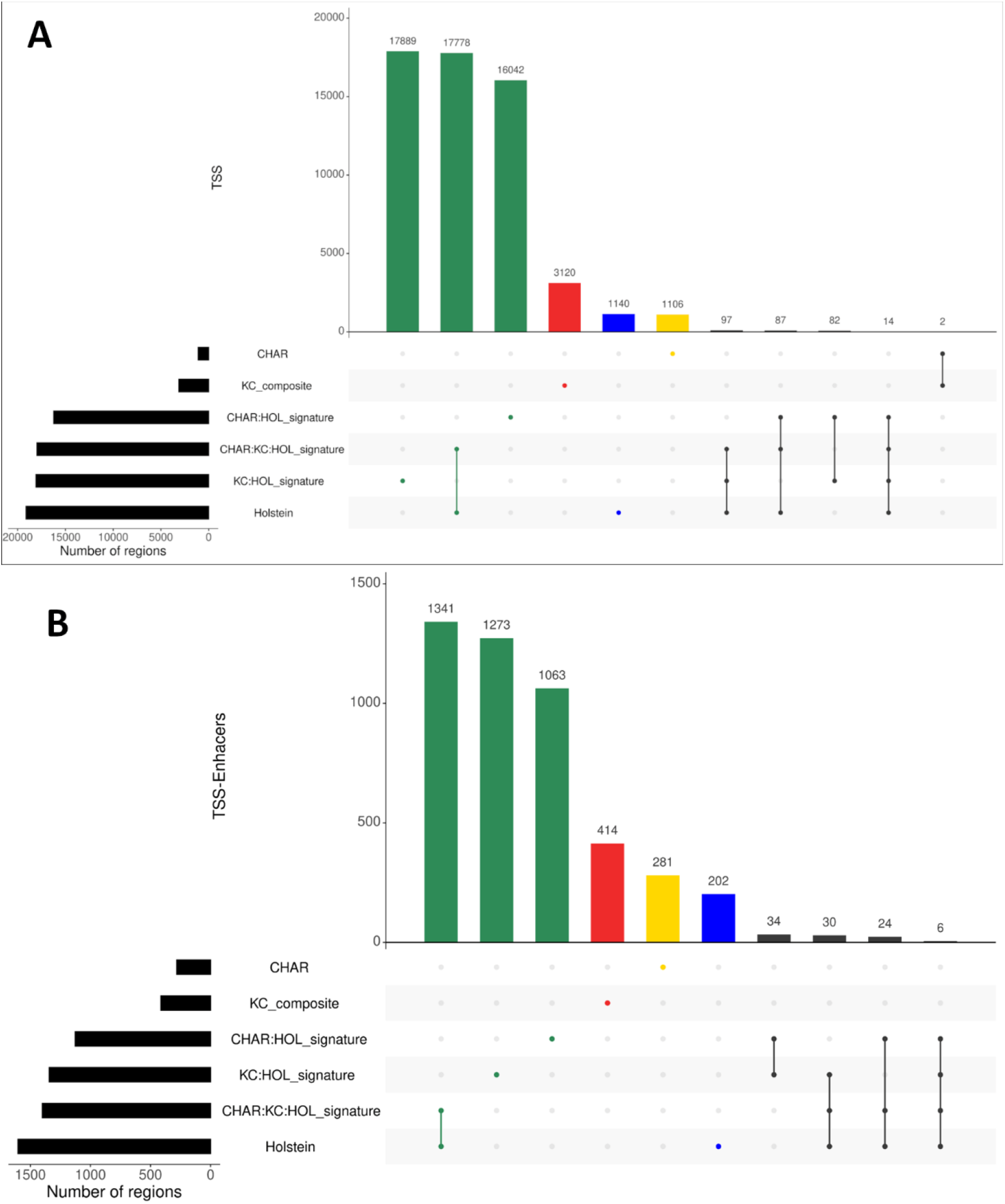
The population-specific analysis of the (A) TSS and (B) TSS-Enhancers regions in 3 populations of cattle. The intersection analysis produced 6 sets of TSS and TSS-Enhancers as following: CHAR regions only present in tissues derived from Charolais x Holstein F2 animals, KC_composite regions only present in tissues derived from Kinsella composite animals, Holstein regions only present in tissues derived from Holstein Friesian animals. CHAR:HOL_signature, KC:HOL_signature were regions shared between the Holstein Friesian dataset and 2 other populations separately. CHAR:KC:HOL_signature a commonly shared set of regions amongst all 3 population of cattle.

### Multi species comparative analysis using the Fantom5 and sheep CAGE datasets

We compared the predicted TSS regions identified within the sheep and cattle CAGE dataset with the previously released Fantom5 CAGE datasets (Bertin et al. 2017). Multi species metrics for these CAGE datasets are shown in Table 3.

**Table 3.** Comparison of the mapped TSS and annotated genes identified in other CAGE datasets (Fantom5, OvineFAANG and BovReg). Column ‘Genes’ corresponds to only the genes that were annotated using the CAGE data (using the 2/3^rd^ tissue representation threshold). The table is sorted (in descending order) by the number of unique TSS identified in each genome.

| Species | Genome | TSS↓ | Genes |
| --- | --- | --- | --- |
| Human | hg38 | 209,911 | 31,184 |
| Mouse | mm10 | 164,672 | 30,501 |
| **Cow** | **ARS-UCD1.2_Btau5.0.1Y\$** | **51,295** | **15,364** |
| Chicken | galGal5 | 32,015 | 7,759 |
| Rat | rn6 | 28,497 | 13,719 |
| **Sheep** | **Oar rambouillet v1.0£** | **28,148** | **13,912** |
| **Sheep** | **ARS-UI_Ramb_v2.0\*** | **27,011** | **13,771** |
| Rhesus monkey | rheMac8 | 25,869 | 8,047 |
| Dog | canFam3 | 23,147 | 5,288 |

\* NCBI RefSeq gff3 annotation v104
\$ Ensembl gff3 annotation v106 track lifted over to 1000bulls reference genome
£ NCBI RefSeq gff3 annotation v100

In this study we have identified the largest number of TSS/promoter activity regions in a non-model organism to date, annotating more than 15,364 genes in the cattle genome. By remapping of the data to the current ARS-UI_Ramb_v2.0 compared to the original reference assembly Oar_rambouillet_v1.0, the number of TSS identified in the sheep CAGE dataset was slightly reduced (∼ 5% less TSS and ∼ 2% less annotated genes). A comparison of the CAGE (TSS) annotated genes from different avian and mammalian species showed high levels of overlap with both cattle and the remapped sheep CAGE datasets. Overall we were able to identify 11,069 genes and their associated TSS unique to the cattle genome (Figure 8).

**Figure 8.**
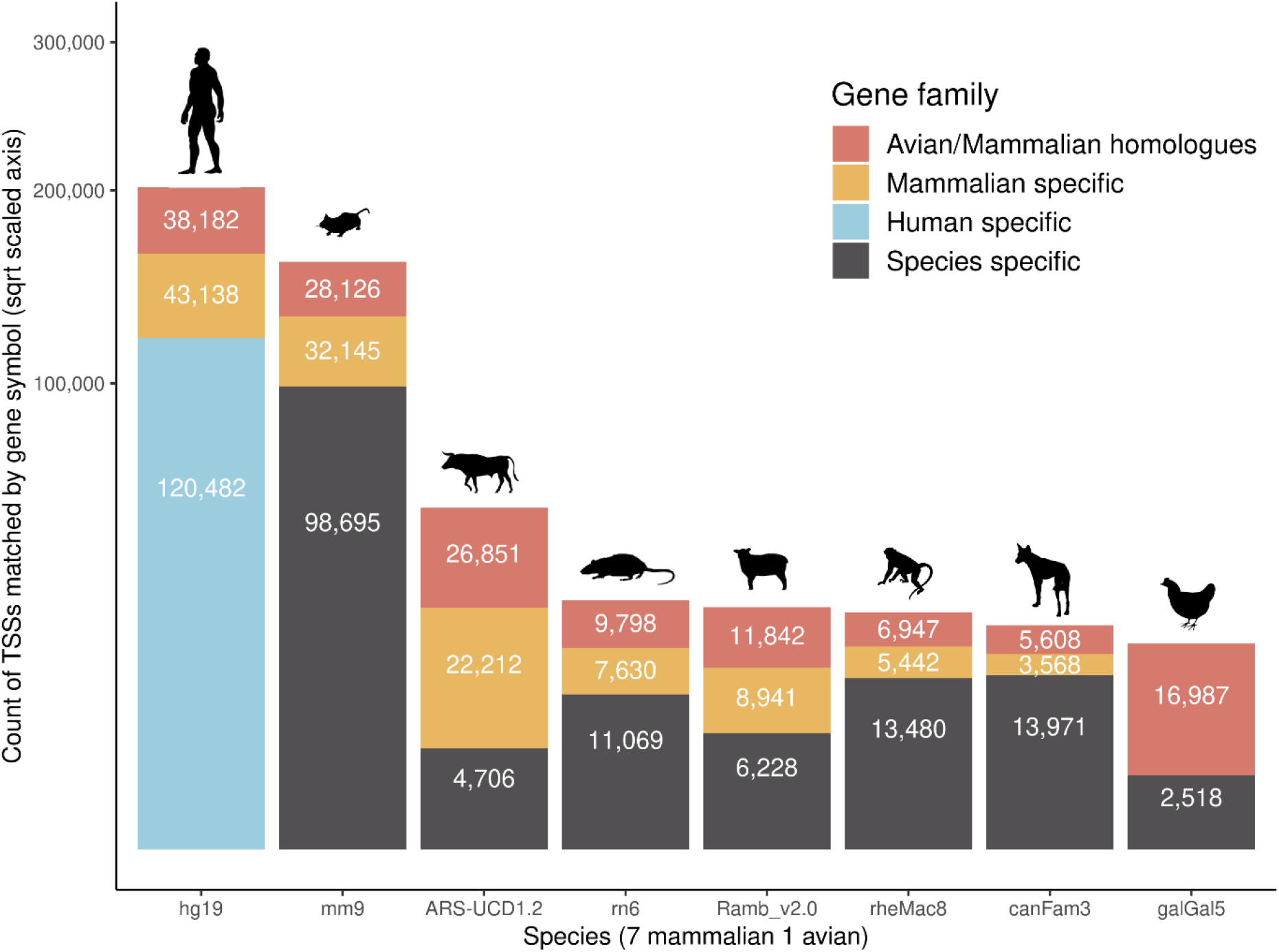
Distribution of the annotated TSS regions (gene symbols) across 8 species. The Fantom5 human, mouse, rat, dog, rhesus monkey and chicken CAGE predicted promotor regions were analysed and compared with the cattle and sheep annotated datasets. The TSS regions, annotated by gene symbols, were coloured in each dataset based on Avian/mammalian origin (gene symbols present in all 8 species), Mammalian specific (7 mammalian species), Human or species specific (gene symbol unique to human or each species).

## Discussion

High resolution mapping of the actively transcribed regions of the genome can help to identify the drivers of gene expression, regulation and phenotypic plasticity (Tippens et al. 2018). Defining transcription start sites (TSS) within promoter regions can provide information about how genes controlling traits of interest are expressed and regulated. To improve TSS and enhancer annotation of the current reference genome for cattle (ARS-UCD1.2), we used CAGE sequencing. We identified more than 51k unique putative TSS coordinates (22% un-annotated regions of the cattle genome) compared to 27k TSS in sheep (7% unannotated regions). The promoter plasticity captured by employing sampling of 24 tissue types from 3 divergent cattle populations for this study resulted in identifying multiple TSS per transcript (mean 3.1 median 2) in the cattle dataset with high reproducibility across tissue types (support mean 23.7 median 24; n=24) and samples (support mean 91 median 94; n=102). This dataset provides a high confidence set of promoter annotations for the cattle transcriptome including ‘novel’ promotors not previously annotated in the available NCBI v.106 and Ensembl v.106 annotations (25% of TSS overlapped with currently annotated promoters and were 1<kb proximal to annotated gene models).

Similar to previously reported studies in cattle (Goszczynski et al. 2021), pig (Halstead et al. 2020; Kern et al. 2021) and human (Andersson et al. 2014) we also identified both tissue and population specific sets of TSS and TSS-Enhancers. Recently new genomic resources have been generated for farmed animal species, including pangenomes and breed-specific reference quality assemblies e.g.(Li et al. 2019; Crysnanto et al. 2021; Talenti et al. 2022). Usage of breed specific genome assemblies can provide a more accurate picture of structural variants specific to a population of animals and ensure better mapability for sequence data in reference guided approaches. Identifying breed-, population- or species-specific promoter complexity can help to harness the full potential of these assemblies as tools to inform genomics enabled breeding programmes e.g. reviewed in (Georges et al. 2018; Clark et al. 2020). We identified full tissue support for TSS and TSS-Enhancer regions unique to each of the 3 populations of cattle in this dataset. The highest number of TSS and TSS-Enhancers regions were present in the most diverse population (Kinsella composite). This finding further highlights the value of including samples from more than one breed in creating reference annotation datasets.

Using methodology for identifying longer stretches of super-enhancers (Thodberg et al. 2019) we also identified 16 genomic stretches (the longest of which was 25kb) from 53 candidate bi-directional TSS-Enhancer clusters. The overlay of these super-enhancers that we performed with previously reported copy number variants for the *PCLG1* gene provides a valuable insight to the regulatory landscape for this gene. *PCLG1* has been identified as a stature (chest width) phenotype associated quantitative trait loci (QTL) target in Simmental (dual purpose) cattle by (Doyle et al. 2020). It has also been reported as a differentially expressed gene between high/low gain vs high/low intake amongst n=143 cross-bred steers from 15 different beef breeds by (Zarek et al. 2017). In addition, the expression of *PLCG1* has been shown to be downregulated due to maternal under nutrition in the muscle tissues of Japanese Black calves raised on a low nutritional value diet (Muroya et al. 2021). Given the critical role of *PLCG1* in both muscle growth and metabolism in beef cattle the knowledge of its associated super-enhancer coordinates and co-expressed promoter regions across tissues could serve as a guide for future functional validation, gene editing or marker selection studies. Another CNV associated super-enhancer region identified in our dataset was *TANGO2*, a golgi system associated protein coding gene mainly associated with mitochondrial disease (Heiman et al. 2022). *TANGO2* has been shown to be over-expressed in seminal plasma of lowly/sub fertile bulls (Muhammad Aslam et al. 2014) and is highly associated with multiple heifer fertility traits in the Holstein cattle population (Chen et al. 2021). Knowledge of the regulatory landscape of genes such as *TANGO2* provides a path for understanding the role of these genes in cattle fertility phenotypes.

We also compared the sheep and cattle datasets with other publicly available TSS and TSS Enhancer genomic tracks for mammalian and avian species to further identify promoters specific to the cattle genome. Using a homologue matching approach the TSS annotation of the cattle dataset captured the highest number of mammalian and (or) avian genes families represented in the datasets, after human and mouse, demonstrating how comprehensive the dataset generated for cattle is. Such information could be used to understand how the genome controls traits in different species, and to identify regions that are important for conservation in breeding programmes.

The CAGE data produced for this study when combined with transcriptomic datasets (mRNA, miRNA and total RNA-Seq) produced by BovReg partners will provide a new comprehensive transcriptome annotation for the cattle genome, as a resource for the farmed animal genomics community. These improved promoter annotation (TSS and TSS-Enhancers tracks per tissue type) will also be available to the community using the FAANG data portal Genome Browser at (https://api.faang.org/files/trackhubs/BOVREG_CAGE_EUROFAANG/) upon publication.

## Data availability

The raw sequence data for all the CAGE-Seq libraries is available via the European Nucleotide Archive and the https://data.faang.org (BovReg/EuroFAANG portal) under BioProject ID

PRJEB43235. The tissue level TSS and TSS-Enhancers regions tracks are also available FAANG data portal Genome Browser at (https://api.faang.org/files/trackhubs/BOVREG_CAGE_EUROFAANG/) and <u>FAANG Genome Browser</u>.

## Code availability

The code and documented analysis pipeline developed in NextFlow DSL2 syntax (di Tommaso et al. 2017), is available at https://github.com/mazdax/nf-cage.

## Supplementary materials

All the supplementary files and figures associated with this publication are available at the following link: https://doi.org/10.6084/m9.figshare.21769649

## Ethics statement

The Canadian sampling study was approved by Animal Care and Use Committee at the University of Alberta (AUP00002592). Animals were transported and euthanized according to the NFACC Code of Practice for beef cattle (National Farm Animal Care Council (DCF-NFACC) 2013). Necropsy and tissue collections were performed under site-specific ethics approval by qualified research personnel at University of Alberta Canada (Animal Use Protocol #00002592), University of Liege, Belgium and the Research Institute for Farm Animal Biology, Germany. The Belgian sampling study had local ethical approval (*Commission d’Etique Animale; Dossier* #17-1948) and complied with the relevant national and EU legislation. In Germany, all experimental procedures were carried out according to the German animal care guidelines and were approved and supervised by the relevant authorities of the State Mecklenburg-Vorpommern, Germany (State Office for Agriculture, Food Safety and Fishery; LALLF M-V/ TSD/7221.3–2.1-010/03).

## Authors contribution

MS developed the nf-cage pipeline, analysed the data, produced all the figures and drafted the initial draft of the manuscript. ELC designed the study, co-wrote the manuscript with MS and edited the final version. RC prepared the CAGE-Seq libraries and undertook sequencing. CK designed the experiment and coordinated the sampling/shipment process for the German samples with DB. GP designed the experiment and organised the sampling/shipment process for the Canadian samples. SD, CC and GCMM collected, processed and shipped extracted RNA from all the collected samples and arranged shipment of these to RC. CK coordinates the BovReg project as a whole. CC, EC and GCMM coordinated the transcriptomic analyses for the BovReg project.

## Acknowledgements

We would like to thank Dr. Haruko Takeda, MSc. Lijing Tang, and Miyako Sakai (GIGA, University of Liège, Belgium) for their help in sampling, storage and shipment of the samples. We would also like to thank Dr Tim Regan for his advice and input for the KDE analysis, Dr Jose Antonio Espinosa-Carrasco for NextFlow code development, rechecking and troubleshooting of the nf-cage pipeline. The contribution of the following are acknowledged for their work in collecting tissue samples from the Kinsella Composite animals: Janelle Jiminez and Carolyn Fitzsimmons for the selection of animals, organization of the tissue sampling team, and maintenance of tissue inventories, Leanna Grenwich, Leluo Guan, ChangXi Li, and Manuel Juarez and their staff as well as the facility staff at the Roy Berg Kinsella Research Station and the abattoir staff at the Agriculture and Agri-Food Canada (AAFC) Lacombe Research and Development Centre, AB, Canada, for cattle husbandry and tissue sampling.

